# Andaman mangrove sediments: source of nutrients and sink of heavy metals

**DOI:** 10.1101/431262

**Authors:** A.K. Mishra, K. Manish

## Abstract

Andaman Islands (AI) of India is a biodiversity hotspot of mangroves but biogeochemical dynamics of AI is less understood. We collected sediment samples of four AI mangrove sites and one site without mangroves for nutrients and trace metal analysis. Samples were collected from each site at the inlet of seawater (Zone A) and the other 500m into the mangrove creek (Zone B). Nutrients (sulphate, ammonium, nitrite and nitrate) levels, organic matter (OM) and carbon content were higher at Zone B of mangrove ecosystem due to the higher OM content from mangrove leaf litter decomposition and microbial degradation. Metal (Pb &Cd) content of zones with and without mangroves were similar and Igeo values indicated moderate contamination of mangrove zones of AI due to lack of anthropogenic pollution. Our results suggest mangrove ecosystems of AI are uncontaminated from heavy metals and are source of nutrients to the oligotrophic coastal ecosystems of Andaman Sea.

## Introduction

Mangrove forests are one of the dominant and productive estuarine coastal ecosystems^1^ and are biogeochemically complex and active^2^. Mangrove ecosystem plays a significant role in the nutrient, organic matter and carbon cycling resulting in better functioning of the coastal ecosystem^3,4^ and providing ecosystem services such as nurseries for fish juveniles^5^. On global scale mangroves contribute to >10% of the total export of land derived dissolved organic carbon (DOC) to the coastal oceans, while covering only <0.1% of the continental surface^6,7^. They also help in sequestration and burial of 23% of global organic carbon in coastal oceans^8^. Mangrove ecosystems of the estuarine zone are more dynamic and complex resulting in export of land mediated nutrients to oceans especially in the oligotrophic waters of tropics and subtropics ^9,10,11^ that benefits the adjacent coastal seagrass ecosystems ^12^ or coral reefs^11,7^.

Mangrove ecosystem are known for their transport of organic materials to the nearby coastal ecosystem^13^, where mangrove leaf litter forms the base of detritus food-web and this detritus is converted by the microbial biomass as food source for higher trophic levels ^14,15,16^. Along with detritus transfer mangrove ecosystem also exports dissolved ammonium, silicate and phosphorus ^11,7^ to the adjacent coastal ecosystem. The net export of matter from mangrove ecosystem depends on the various physico-chemical and biological processes within the mangrove sediments and water column^11^. To export materials, mangrove ecosystems act as both source and sink of materials, directly by absorbing dissolved nutrients and carbon from terrestrial run-off^17^ and indirectly absorbing them from the deposited sediment^18^. However, lower oxygen content of mangrove sediments^19^ prevents microbial degradation of matter, resulting in a transfer of undegraded organic matter to adjacent coastal ecosystem that can be utilized by other organisms in oxygen rich environment^20^. This magnitude of transfer of materials to adjacent coastal ecosystems are dependent on the amount of fresh water input into mangrove creeks^21^. Though mangrove ecosystems are highly productive and have a rich source of organic matter, they are generally nitrogen and phosphorus deficient ^22,23^. To overcome this nutrient deficiency, mangrove ecosystems depend on the efficient internal nutrient cycling of organic matter by microbial (bacteria and fungi) biomass in the sediments ^22,23^ and mangrove leaf litter plays an important role in providing 40% water soluble components for bacterial biomass ^6^.

Trace metals in marine sediments are particularly of high importance due to their continuous persistence in the sediments resulting in bioaccumulation and potential toxicity to the sediment associated biota^24,25,26^. Mangrove ecosystem being at the forefront of terrestrial and marine aquatic environments acts as a trapping zone of terrestrial runoff and organic material debris. As the organic matter (leaf litter and wood) decays it leaches out various elements and land runoff also acts as source of various pollutants that gets concentrated in the sediment resulting in mangrove ecosystem acting as a sink for metals^27,28^. These sediments act as potential secondary source of trace elements that can be released back into the water column under change in environmental conditions such as temperature and pH ^29,30^and under various disturbances such as bioturbation, resuspension etc. These metals in water column and sediments becomes bio-available and are transferred into higher trophic levels^31,27^ exerting possible toxic effects on the associated biota through biomagnification^32,27^. However, in mangrove ecosystem the trace metal content in the water column is lower than sediment due to daily tidal variation and low residence time^33^, which increases the importance of mangrove sediments as proxies of the environmental metal contamination. Higher metal contamination due to various anthropogenic influences have been observed at different mangrove ecosystems of India, for example Lead contamination in Pichavaram mangrove sediments of Tamilnadu, India^34^, Copper, Manganese, Nickel and Zinc contamination was observed in the mangrove sediments of Sundarbans delta, India ^33^.

In Andaman and Nicobar Islands (ANI) the coastal areas are enriched with abundant mangrove forests covering 644 square kilometre^35,36,37^ due to favourable environmental conditions, such as heavy rainfall, short dry season and high tidal fluctuation^38^. Mangrove ecosystems of India represent 3% of world mangrove ecosystems ^35^ and ANI represent 13% of the total mangrove ecosystems of India^39,35^ and 50% of the global mangrove species^40^. Though these islands are enriched with mangrove forests, most of the studies on mangroves have been focused on biodiversity assessment^40,41^ ^42^ distributions and vegetative structure^43,41^ or valuation of mangrove ecosystem services^44^. Very few studies have been carried out on the nutrient concentration of mangrove sediments of Andamans, such as nitrite, nitrate and phosphate levels in the sediments of Chidiyatappu and Rangat Bay for phytoplankton productivity estimates of mangrove ecosystem^42^,whereas nitrite, nitrate, ammonium and phosphate concentrations of mangrove sediments and water column of Aerial Bay and Chidiyatappu Bay were reported for determining water quality of mangrove ecosystems^37,45^. Similarly, nutrients such as nitrite, nitrate, ammonium and phosphate concentration of Aerial Bay and Rangat Bay with mangroves was reported for water quality monitoring ^46^. Other studies have focused on the influence of anthropogenic activities on coastal ecosystem of these islands, such as influence of sewage and land run-off on the microbial biota and nutrient concentrations of Portblair Bay^47^, changes in bacterial and physico-chemical properties of seawater due to land run-off^48^, whereas hypoxic conditions created by sewage disposal due to human activities have been reported near mangrove ecosystem of Phoenix Jetty, South Andamans ^49^.

Saying that, the waters of Andaman Sea are oligotrophic^49^, which requires the input of necessary materials from the adjacent coastal mangroves and seagrass ecosystems for better ecosystem functioning. This increases the importance of mangroves in production, cycling and transfer of nutrients to adjacent coastal ecosystems. So, the objective of our work is to quantify the various nutrient, organic matter, carbon content in four mangrove ecosystems of Andamans and compare the concentrations with one site without mangroves to understand the influence of mangrove ecosystem in nutrient generation and transfer. Five sites will be used and from each site a zone near the seawater inlet will be compared with a zone inside the mangrove ecosystem to determine the transfer of nutrients from mangrove ecosystem. Trace metal levels will be assessed to understand the metal pollution levels of these mangrove ecosystems.

## Material and Methods

### Study sites

The study area comprises of five different sites of Andaman Islands (AI) in Andaman Sea (Fig.1). Andaman and Nicobar Islands (ANI) are situated in the South-east coast of Indian subcontinent in Bay of Bengal, Indian ocean. ANI are surrounded by Andaman Sea and harbouring a rich diversity of flora and faunal^36,37^. Out of these five sites four were situated in mangrove ecosystem (Carbyn cove, Chidiyatappu, Burmanala and Kalapathar) and one was without mangroves (Marina Jetty). All the five sites were exposed twice daily at low tides. At all four sites of mangrove ecosystem, one area was selected close to the inlet of seawater (zone A) and the second one was at 500m upstream into the mangrove zonation (zone B). Similar zonation was used for Marina Jetty with no mangroves. The abundant mangrove population at Carbyn cove, Chidayatapu and Kalapathar were *Rhizophora* species, whereas Burmanala was dominated by *Bruguiera* species, however at four sites both species were present.

**Fig. 1.**
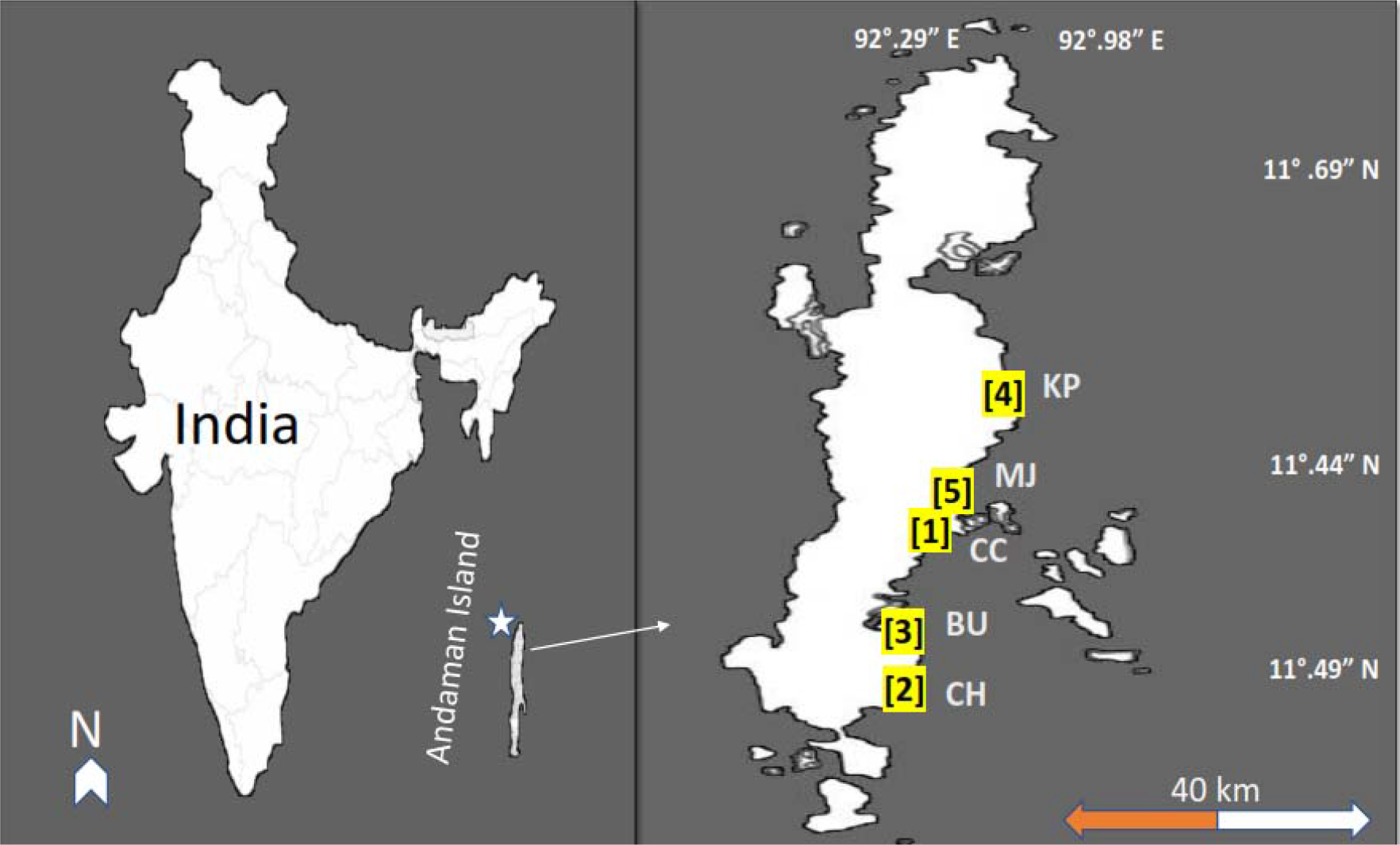
Five study sites of Andaman Islands in Andaman Sea, India. [1] Carbyn cove (CC) [2] Chidayatapu (CH) [3] Burmanala (BU) [4] Kalapathar (KP) [5] Marina Jetty (MJ).

### Sampling

Sampling was carried out in the pre-monsoon (April-May) season of ANI. At all five sites from Zone A and Zone B three points of 1m apart were selected and from each point sediment samples (n=3×2) were collected during low tide from a depth of 05 cm with plastic syringes with their tip cut off. The syringes helped to keep the sediment intact and easy to suck up in the syringe. Collected sediment samples were stored in plastic bags, stored in dark boxes and transported to laboratory for further analysis. Salinity, pH and temperature of the sediment samples were measured in the field using a Salinometer (SR-028, Aichose) and Hydrolab (Hydrolab Quanta, OTT, Hydromet) respectively. In the laboratory sediment samples were placed on big tray on white paper and air dried in room temperature. The dried samples were kept in plastic bags till further analysis.

Nutrient (sulphate, inorganic phosphate, ammonia-nitrogen, nitrite-nitrogen, nitrate-nitrogen, organic matter and carbon content) were analysed using UV-Visible spectrophotometer (Perkin Elmer, Lambda 35) using the methods described in Coastal Ocean Monitoring and Prediction System for Indian coast ^50,51^. All samples for nutrients were analysed in triplicates and quality control procedures ^50,51^ were followed by careful standardization and blank measurements.

### Metal analysis ^51^

Dried sediment of one gram was taken in a flask to which 20 ml of analytical grade triacid (HNO_3_: H_2_SO_4_: HCLO_4_: 14:1.5:4.5) was added and heated in a chamber till complete digestion of the sediment samples. Following the digestion, the sample was allowed to cooldown and then it was filtered to a 100-ml volumetric flask, distilled water was used to make the volume up to 100 ml of the filtrate. These samples were then used for the analysis of heavy metals using atomic absorption spectrophotometer (AAS). For Lead (Pb) the wavelength used in AAS was 283.31 nm while that for Cadmium (Cd) was 28.80 nm. Distilled water was used as blanks and samples and standards ^51^ were analysed as triplicates.

### Geo-accumulation index^52^

To assess the contamination level of sediments Geo-accumulation index was used. The index of Geo-accumulation (I_geo_) was computed using the following equation

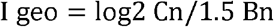

Where C_n_ is the measured concentration of the element in the sediment, B_n_ is the geochemical background value derived from average shale value of the element in earth’s crust^53^ and the constant 1.5 represents natural fluctuations for the element concentration in the environment with very small anthropogenic influence. The results obtained were compared to the below presented six classes of the geochemical index ^52,54^.

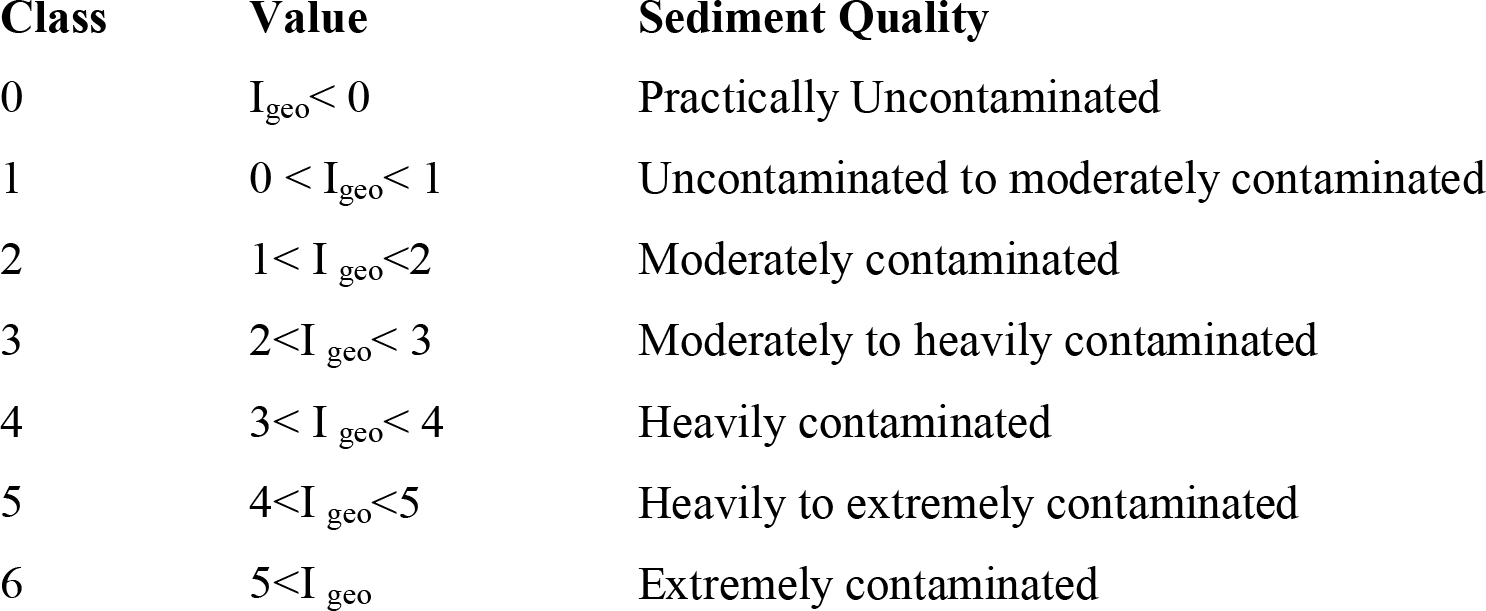

### Data analysis

All data was pre-checked for normality and equal variance. A two-way ANOVA using site (five sites) and location (Zone A and Zone B) as factors were used to test the significant difference (p<0.05) between variables. Data that did not show significant difference were log transformed and tested again. After ANOVA, Holm-Sidak pairwise multiple comparison was used to check the significant differences among Sites and locations using SIGMAPLOT statistical software^55^. All values in text are expressed as mean± standard error. Based on nutrients and heavy metal contents a hierarchical clustering (Bray-Curtis Similarity) was applied to describe the similarity between all five Sites using PRIMER v.7 software ^56^.

## Results

The physical parameters were similar among the mangrove and without mangrove sediments. The pH ranged between 7.96±0.03 to 7.97±0.05 among the mangrove sites, whereas at sites without mangrove it ranged between 7.95±0.03 to 7.96±0.02. Salinity was similar among all sites ranging from 31.48±0.02 to 31.50±0.03. Temperature range at site with mangroves 30.63 to 30.66°C, whereas with sites without mangroves it ranged from 31.45 to 31.55°C.

Nutrient concentrations of sediment at the five sites of AI were significantly different near the inlet of seawater (Zone A) and within mangrove ecosystem (Zone B). The four sites (Carbyn cove, Chidiyatappu, Burmanala and Kalapathar) with mangroves were observed with higher sulphate levels in sediment than Marina Jetty without mangroves. However, phosphate levels were higher at sites without mangrove compared to sites with mangrove ecosystem (Fig.2). The sediments at zone B of Carbyn cove (2.55± 0.02 µg/g) were observed with the highest sulphate levels followed by Burmanala (2.21± 0.01 μg/g) and Kalapathar (2.13± 0.03 μg/g). Sulphate levels at zone B of Carbyn cove was 1.8-fold higher than zone A (Fig.2.A). Phosphate levels was highest at zone A of Chidayatapu (1.97± 0.10 μg/g) followed by Marina Jetty (1.17± 0.01 μg/g) and Kalapathar (1.01μg/g ± 0.10), exceptions were Carbyn cove and Burmanala. Phosphate levels of Burmanala at zone B was 1.5-fold higher than zone A (Fig.2.B).

**Fig. 2.**
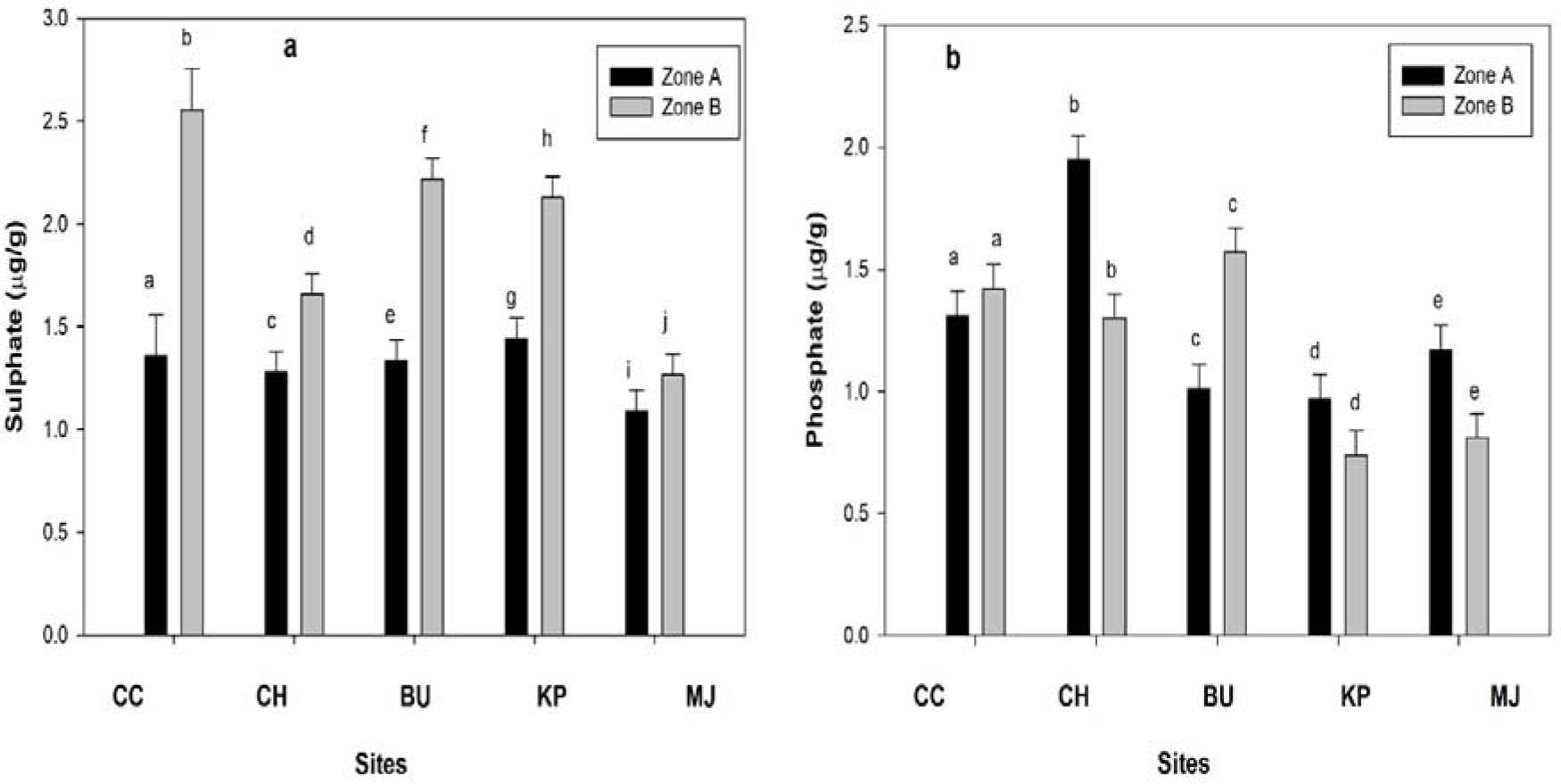
Sulphate (a) and phosphate (b) levels in sediment of mangrove ecosystem at five sites of Andaman Islands. Error bars represent standard errors. Significant difference between zones are indicated by different letters. Carbyn cove (CC), Chidayatapu (CH), Burmanala (BU), Kalapathar (KP) and Marina Jetty (MJ).

Ammonium and nitrite concentration were higher at zone B of all sites, whereas for nitrate, Kalapathar and Marina Jetty were observed with higher concentrations at zone B (Fig.3). Ammonium concentration at zone B of Carbyn cove (1.31± 0.10 μg/g) and Marina Jetty (0.30 ±0.01 μg/g) was 3.6-fold and 3.4-fold higher than their zone A respectively (Fig.3A). Nitrite concentration of Carbyn cove (0.34 ± 0.01 μg/g) was highest followed by Kalapathar (0.26 ±0.01 μg/g) and Marina Jetty (0.19 ±0.01 μg/g). Nitrite concentration at zone B was 2.2-fold higher than zone A of Carbyn cove (Fig.3 B). Nitrate concentration was higher at Chidayatapu zone A (0.60 ±0.01 μg/g) followed by Burmanala (0.32 ±0.01 μg/g) and Kalapathar (0.27 ± 0.01 μg/g), though the nitrate concentration of zone B of Marina Jetty was 12.4-fold higher than zone A (Fig.3C).

**Fig. 3.**
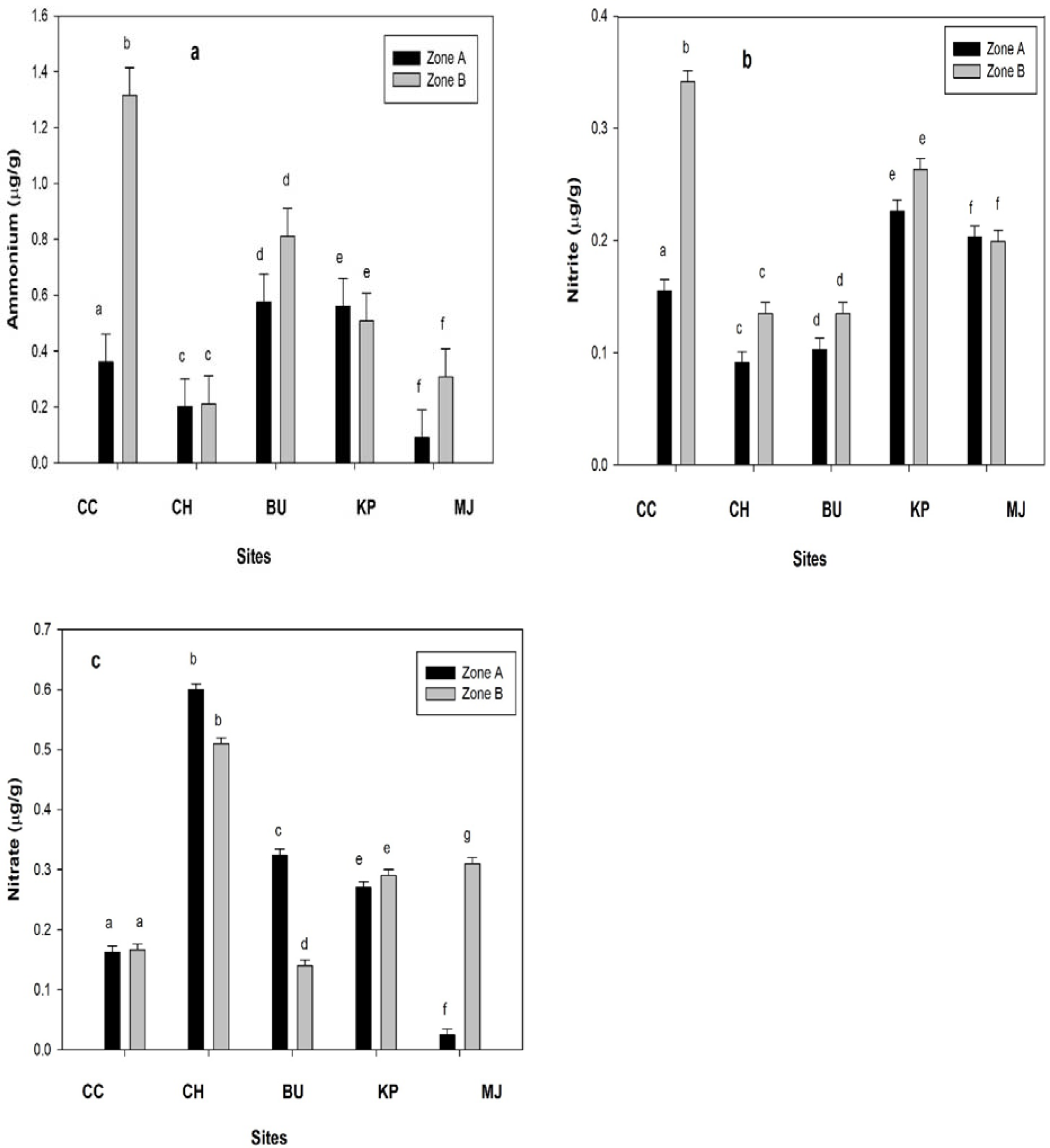
Ammonium (a), nitrite (b) and nitrate(c) levels in sediment of mangrove ecosystem at five sites of Andaman Islands. Error bars represent standard errors. Significant difference between zones are indicated by different letters. Carbyn cove (CC), Chidayatapu (CH), Burmanala (BU), Kalapathar (KP) and Marina Jetty (MJ).

Sediments within mangrove zones were observed with higher organic matter and carbon content than zones without mangroves. Among all five sites organic matter content was highest at Chidiyatappu (6.24±0.20) followed by Kalapathar (5.62 ±0.20) and Carbyn cove (5.99±0.20) zone B (Fig.4a). Carbon content was highest at zone B of Burmanala (3.80 ± 0.10) followed by Chidiyatappu (3.62±0.20) and Carbyn cove (3.44±0.20) (Fig.4b). Carbyn cove and Kalapathar zone B organic matter and carbon content was 4.2-fold and 1.5-fold higher than their zone A respectively. However, at Marine Jetty zone A was observed with higher organic matter and carbon content than Zone B (Fig.4).

**Fig. 4.**
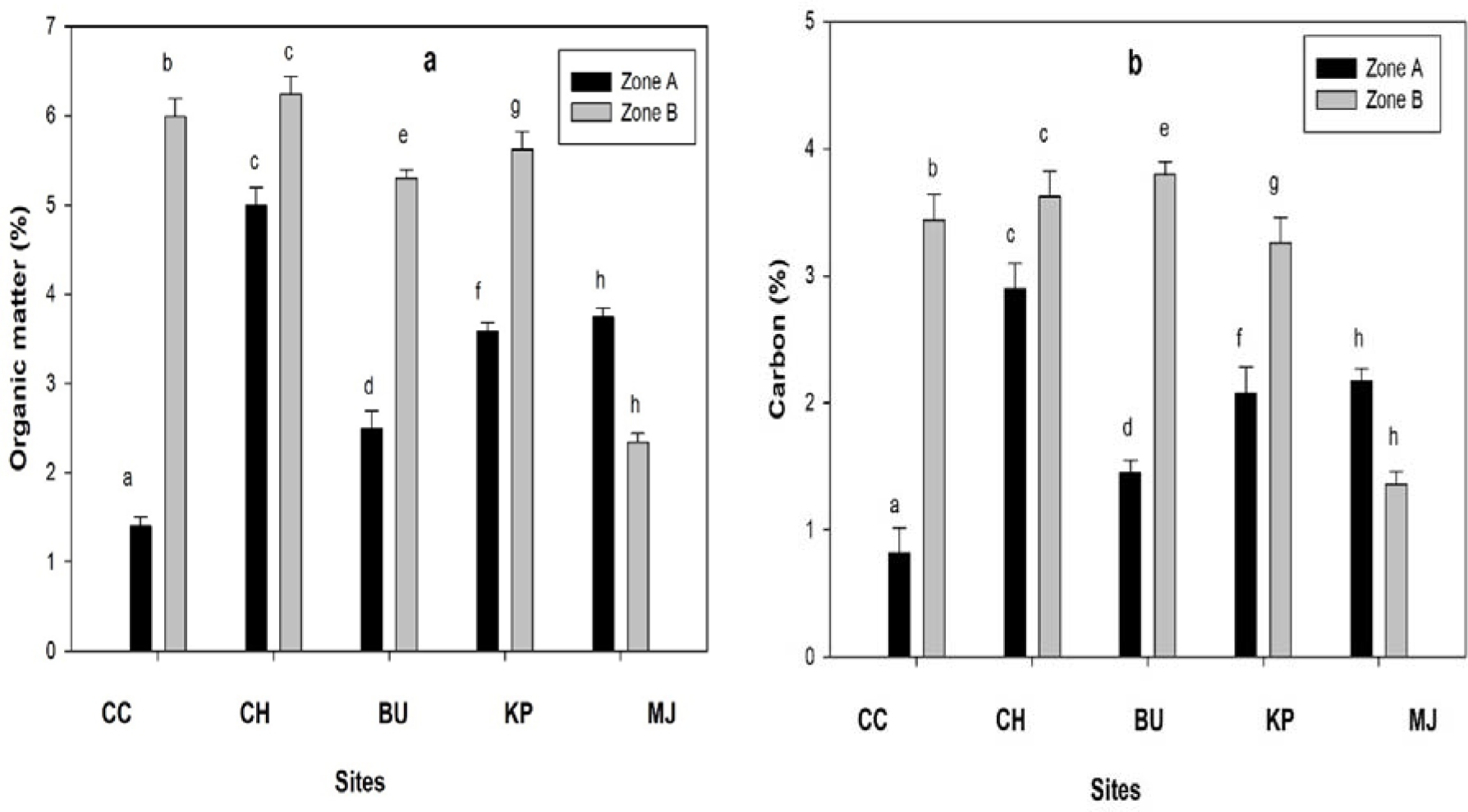
Organic matter (a) and carbon (b) content in sediment of mangrove ecosystem at five sites of Andaman Islands. Significant difference between zones are indicated by different letters. Carbyn cove (CC), Chidayatapu (CH), Burmanala (BU), Kalapathar (KP) and Marina Jetty (MJ).

Metals (Cd and Pb) concentration were different and significant for sites with and without mangroves. Cd concentration was higher within the mangrove sediments at zone B of Carbyn cove, Chidiyatappu and Kalapathar than zone A, exception was Burmanala and Marine Jetty. However, Cd concentration at zone B of Chidiyatappu (0.30± 0.01mg/Kg) was highest followed by Kalapathar (0.27± 0.01 mg/Kg) and Marine Jetty (0.29± 0.01 mg/Kg). Cd concentration of Kalapathar and Carbyn cove zone B was 1.7-fold and 1.2-fold higher than zone A respectively, whereas Burmanala zone B was 0.6-fold lower (Fig.5a). Pb concentration was higher at zone A of all sites, exception was Kalapathar (Fig.5b). Highest Pb concentration were observed for zone A of Marina Jetty (1.64±0.10 mg/Kg) followed by Chidiyatappu (1.54±0.10 mg/Kg) and Carbyn cove (1.29±0.10 mg/Kg). However, Pb concentration of Kalapathar zone B was 1.14-fold higher than zone A (Fig.5b)

**Fig. 5.**
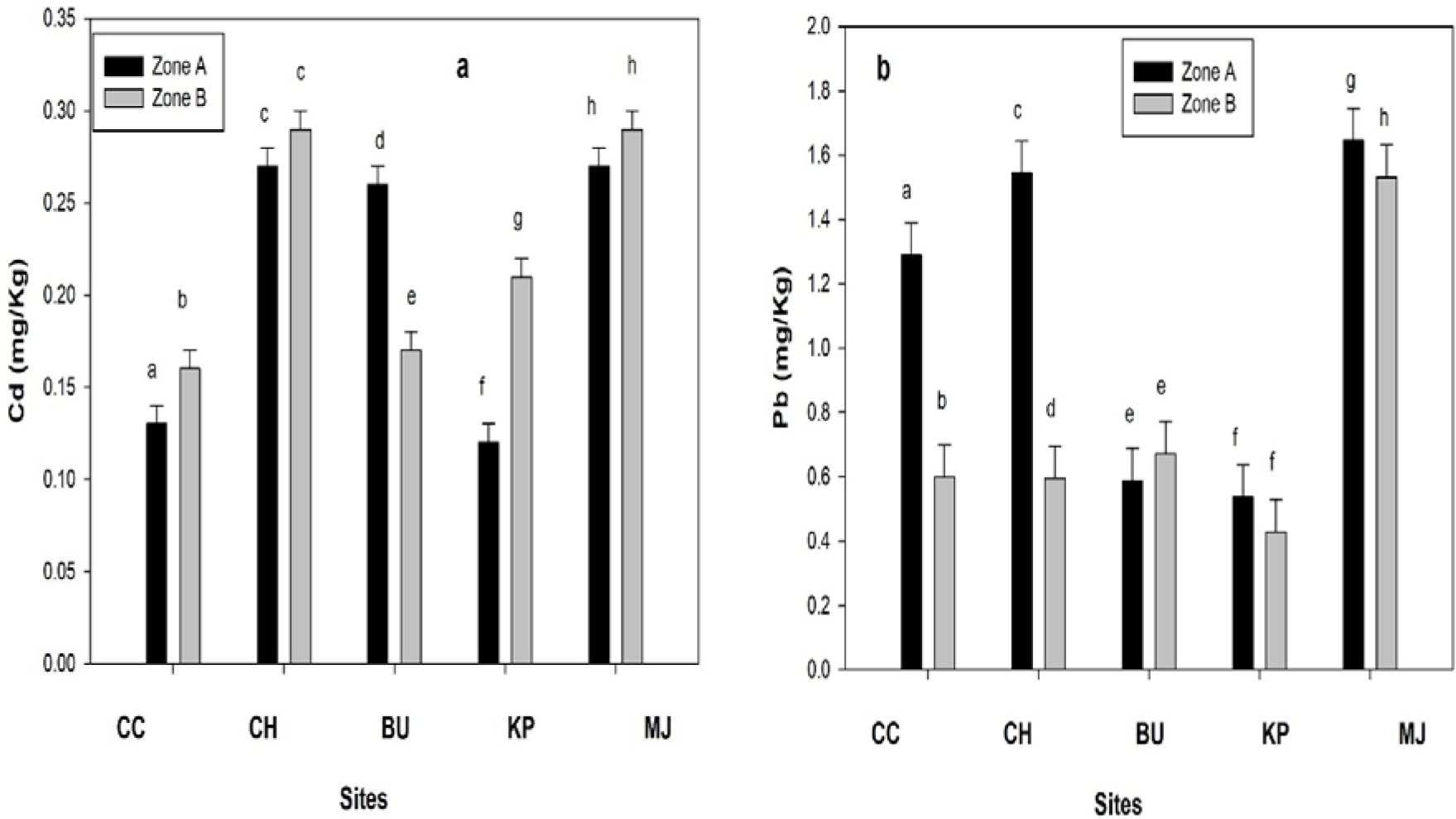
Trace metals Cd (a) and Pb (b) levels in sediment of mangrove ecosystem at five sites of Andaman Islands. Error bars represent standard errors. Significant difference between zones are indicated by different letters. Carbyn cove (CC), Chidayatapu (CH), Burmanala (BU), Kalapathar (KP) and Marina Jetty (MJ).

Igeo values (Igeo<1) indicated uncontaminated to moderate contamination of sediments at all sites except Kalapathar zone B. Cadmium contamination was not observed at all five sites. However, within all five sites, Marina Jetty zone B was observed with higher Pb contamination followed by Chidiyatappu zone A. Overall the mangrove water outlet close to the sea were observed with higher contamination than sediments within mangrove forests (Table 1).

**Table 1.** Igeo values of sediments for Cadmium (Cd) and lead (Pb) at all five sites of Andaman Isalnds Carbyn cove (CC), Chidayatapu (CH), Burmanala (BU), Kalapathar (KP) and Marina Jetty (MJ)

| **Element** | **Sites** | **Zone A** | **Zone B** |
| --- | --- | --- | --- |
| Cd | CC | 0.03 | 0.00 |
|  | CH | 0.05 | 0.01 |
|  | BU | 0.05 | 0.01 |
|  | KP | 0.02 | 0.00 |
|  | MJ | 0.05 | 0.01 |
| Pb | CC | 0.36 | 0.20 |
|  | CH | 0.43 | 0.24 |
|  | BU | 0.16 | 0.10 |
|  | KP | 0.15 | 0.06 |
|  | MJ | 0.46 | 0.66 |

Dendrograms were generated based on the nutrients and trace metal levels of zone A and B at all five sites. Two groups were generated based on 90% similarity index. The first group included zone A and B of Carbyn cove and Chidiyatappu and Burmanala zone A, whereas the second group included rest of the sites. However, zone B of Burmanala and Zone A of Kalapathar were similar with Marina Jetty zone B and A respectively (Fig.6).

**Fig. 6.**
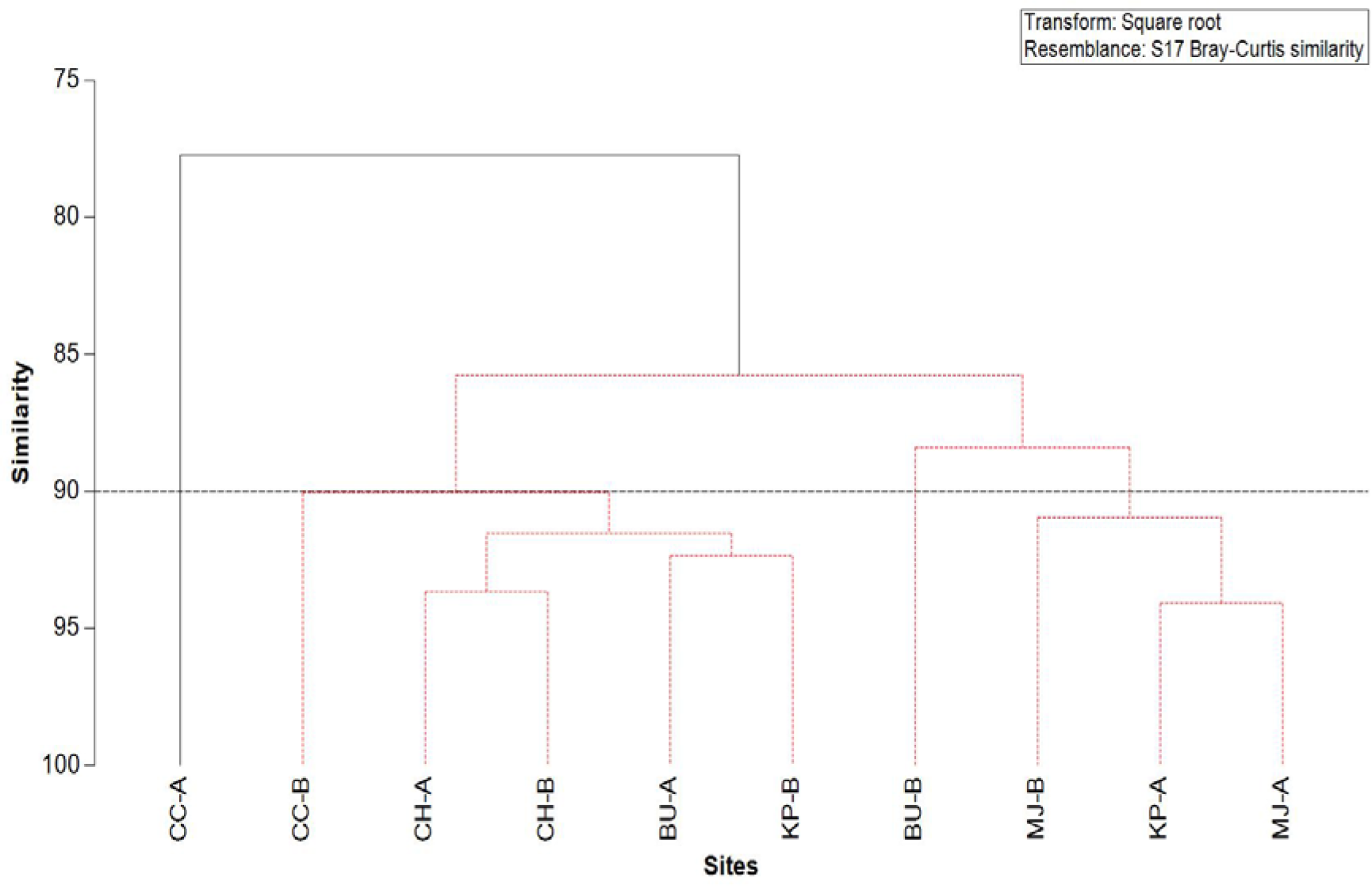
Dendrogram for hierarchical clustering of five sites using group average linking of Bray-Curtis similarities calculated on square root transformed data. The two groups produced by threshold similarity of 90% are shown. Carbyn cove (CC), Chidayatapu (CH), Burmanala (BU), Kalapathar (KP) and Marina Jetty (MJ).

## Discussion

Mangrove ecosystem act as an important source of nutrients, organic matter and heavy metals for the adjacent coastal ecosystems and our results indicate coastal mangrove ecosystems of Andaman and Nicobar Islands abide to this ecosystem function with higher nutrient and lower metal concentration observed within the mangrove sediments. The physical parameters of temperature, salinity and pH observed in our site were like the results observed for the mangrove ecosystems in and around AI for pre-monsoon season ^42,57,45,46,37^.

The sulfate levels observed in our results within the mangrove zone B were higher, as compared to the zones without mangroves which indicates the sulfate richness of mangrove^58^. The range of sulfate levels in the four-mangrove zone A were similar and zone B were 1.3-1.8-fold higher than sulfate levels of Portblair Bay^57^ of AI. For Marine Jetty the range of sulfate (Fig.2) were 1.2-1.4-fold higher than Hado Harbour of AI^57^. Higher sulfate levels within mangrove sediments are result of increased sulfate reduction in the mangrove sediments sourced from decomposed organic matter^3^, facilitated by the sulfate reducing microbial communities found in the upper 4-6cm of the sediment^60^. However, low utilization of sulfate by the dominant microbial community and sulfate not being an essential nutrient also leads to higher levels in sediment. Lower levels of sulfate at zone A than zone B indicates about the differential production of sulfate and the exposure of these zones to wave actions, where zone A receives higher waves, tidal influx, lower resident time and drainage of tidal water compared to zone B within mangroves^61^.

Phosphate levels within the four-mangroves zone B (Fig.2) in our results were higher than Chidiyatappu (1.6-fold)^42^, Sippighat (3.1-fold)^49^and Aerial Bay (4.1-fold)^46^ and were similar to Sundarbans mangroves sediment^61^. Higher phosphate levels within mangrove sediment are due to the land run-off from the nearby agricultural fields, as these mangrove areas are considered pristine in terms of anthropogenic pollution source ^49^. Secondly, the higher organic matter input from the mangrove leaf litter and the microbial degradation, owing to rapid mobilization and recycling of nutrients during decomposition^14,15,16^ also leads to higher phosphate levels. Saying that, the daily tidal influx of decomposed materials to the mangrove ecosystems, their microbial decomposition and mineralization in the sediment also contributes for phosphate generation^15,16^. Although, the mangroves of Andamans lack riverine input during pre-monsoon season the biogeochemical phosphorous cycling seems effective in maintaining the global phosphorous range (<40- <2 μg/g)^22^ for mangrove ecosystem, as observed in our results. Lower levels of phosphate at zone B than at zone A within the mangrove ecosystem suggests the outwelling of phosphate by daily tidal influx to the adjacent coastal ecosystem. Phosphate levels of Marine Jetty (Fig.2) in our study were 1.6-fold and 3.6-fold higher than Phoenix Jetty ^49^ and Aerial Bay^45,46^ respectively and were similar to Rangat Bay^42,46^ which can be related to the continuous input of anthropogenic waste water discharge at Marine Jetty as there are no source of agricultural input at this site.

Ammonium and nitrate levels of Chidiyatappu zone A & B (Fig.3a & c) were 3-fold and 1.2-fold higher than previously observed for Chidiyatappu Bay^37,42^ and Aerial Bay^45^ respectively, whereas ammonium levels of Burmanala zone B were 2-fold higher than previously observed values around AI^37,45,46^ and nitrate levels were 2-fold lower than Rangat Bay^46^ and Sippighat mangrove ecosystem^49^. Ammonium levels of Burmanala and Kalapathar zone A were similar to Aerial Bay^45,46^ and Carbyn cove was similar to Rangat Bay^46^. Marina Jetty ammonium levels were 1.1-fold lower than previously recorded for Aerial Bay Jetty ^45^. Marine Jetty nitrate levels were 2.1-fold lower than Phoenix Jetty^49^ as Phoenix Jetty receives heavy nutrient loading from anthropogenic sources of waste water discharge^61,49^.

Nitrite levels in sediments of mangrove zone B (Fig.3b) of our results were 4.3-fold higher than previously observed results from Chidiyatappu Bay ^37,42^, 3.4-fold than Rangat Bay, were similar with Aerial Bay^45,46^ and were 0.8-fold lower than Rangat Bay^46^ of AI, whereas zone A levels were similar to these sites. Nitrite concentration of Marine Jetty were similar to nitrite concentrations of Aerial Bay^46^ and Hado Harbour^57^.

Higher ammonium levels observed in the surface sediment compared to nitrite and nitrate indicates higher denitrification and anammox activity in the surface sediment of these mangrove ecosystems^61^. The conservation of available nitrate levels within mangrove areas due to limited riverine inputs during pre-monsoon and conversion of these nitrate into ammonium through dissimilatory nitrate reduction also adds ammonium into the surface sediment^62,63^. However, a significant source of nitrate to produce ammonium are generally found in the sediment pore water of pristine mangrove forests like Andaman, which results in intrinsic conversion of nitrate to ammonium^63,45^. Our study indicate, higher nitrate levels within mangrove sediment can be used to produce ammonium. Saying that, ammonium gets easily adsorbed to clay particles, which makes it unavailable to biological uptake, thus leading to higher levels in sediments^64^.

High organic matter content in mangrove sediment as observed in our study also leads to increased microbial anaerobic fermentation resulting in ammonia formation ^65,57^. On the other hand, Marine Jetty without mangroves have lower ammonium levels as the sediments are low in organic substrate (mangrove leaves) resulting in low denitrifying microbial population. The mean organic matter content of zone A and Zone B of four mangrove sites were 2 and 3-fold higher than Korampallam mangrove creek^67^. Higher organic carbon in mangrove sediments are due to decomposition of mangrove leaf litter by microbial activity and hydrolysis of tannins ^58^ to process nutrients^6^. Secondly, these bacterial population are mostly attached to the sediments, where they naturally die and lyse to form dissolved materials^22^ adding organic matter to sediment. This process also prevents transfer of nutrients and dissolved organic carbon between sediment interstitial waters and water column of the mangrove ecosystem resulting in lower nutrient and organic matter content in water column^15^. Mangrove ecosystems are always rich in organic carbon and higher carbon content in our results were similar to the findings at mangroves of Sundarbans^33^.

However, it seems that in the mangrove ecosystems of AI, much of nutrients are always stored in the mangrove generated biomass where microbial decomposition and recycling maintains the required nutrient levels. Nutrient levels in our study (pre-monsoon) were lower when compared to monsoon and post-monsoon levels ^37,57^ indicating the land run off due to monsoon brings heavy load of nutrients from allochthonous origin into the mangrove ecosystem of AI^66^.

The concentration of Cd levels in our study for both Zone A and B were similar to Ennore creek and The Gulf of Mannar^68^ south east coast of India and were 0.5-fold lower than Cd values observed for surface sediments of AI^69^ and various other mangrove locations of Andamans^69^, South east coast of India^70^, Pichavaram Estuary^71^ and Tuticorin coast^67^. Higher Cd concentration at mangrove zone B than zone A can be due to the use of phosphate fertilizers with Cd impurities from the nearby villages that ends up in the mangrove sediments during land run off ^72,73^. Similar higher Cd concentrations in mangrove sediments were observed at Beibu Bay in China Sea^75^. However, higher Cd concentration at Marine Jetty can be due to the urban waste water discharge of Portblair city^76^.

Pb concentration observed in our study were 9-fold lower than Pb levels observed for various other mangrove locations of Andamans^69^, South east coast of India^70^, Pichavaram Estuary^71^, The Gulf of Mannar ^68^, and Tuticorin estuary ^67^. Lower levels of Pb at Zone B within mangrove ecosystem can be a result of high rate of Pb accumulation in mangrove roots followed by low rate of mobilization^27,76^. This kind of storage of Pb in mangrove roots have been observed in *Rhizophora* and *Bruigerra* species at Bhitarkanika, Odisha on the east coast of India^77^ and our sites were observed with same species of mangroves. Another reason for low Pb content in our results can be due to weak correlation of Pb with organic carbon in surface sediments^78^. However, the source of Pb pollution are generally anthropogenic^75,78^ and these mangrove ecosystems of AI are still considered pristine in terms of anthropogenic pollution^69^. As mangrove sediments act as a sink of metals, the geochemical fraction of Pb that originates from rocks around Andamans and are incorporated into mangrove ecosystem due to land run-off and are reflected in our study.

Igeo values for Pb in our study were similar to sediments of south east coast of India^70^ and were 2-fold lower than observed for mangrove sediments of Tamilnadu ^34^, Korampallam creek^67^, which indicates the mangrove sediments of Andamans are less contaminated from anthropogenic pollution than the mainland Indian east coast.

Similarities between the mangrove sites indicates that nutrient and trace metal dynamics of these sites were not much different due to the pristine nature of AI along with low influence of the local environmental factors such as waste water discharge and agricultural run-off ^72,73^. The similarities of Marine Jetty with Burmanala and Kalapathar indicates the influence of higher urban run-off on the nutrient and metal levels of Marine Jetty. Secondly as the samples are collected during pre-monsoon season, mangrove sites have received less land run-off ^66^ providing land derived DOC nutrient generation and trace metals.

Overall our study reflects the low to unpolluted conditions of Andaman Islands by heavy metals as compared to other mangroves zones of India^69,67^. Low pollution in the mangrove locations in our study are due to lack of significant contribution from large scale industries in the AI^69^. The present observed concentrations of Cd and Pb can be due to natural factors like land run-off or erosion from local mineralogy^79^ and deposition of contaminated sediments from the aftermath of 2004 Tsunami^80,56,37^. However due to high anoxic reducing soil conditions, high decomposing activity^81^ and high absorptive capacity of mangrove sediments the metal bound to the sediment fractions may not be bio-available to the plants and the associated biota^82^.

## Conclusion

Our results suggest that mangrove sediments of AI are source of nutrients and heavy metals which are transferred to the adjacent coastal ecosystems through daily tidal influx. Mangroves of AI take part in the generation of organic carbon and nutrients though being in the adverse tropical conditions. Though these mangrove ecosystems of AI are pristine in terms of heavy metal pollution, a constant monitoring is required to look at the effects of anthropogenic pollution. As there is lack of data for sites like Carbyn Cove, Burmanala and Kalapathar, these results can be used as an initial baseline for nutrient and metal concentration of these sites. Role of mangrove ecosystem in generation and transfer of nutrients was evident from our study. So, conservation of mangrove ecosystems around the AI should be a priority to avail the various ecosystem services mangroves provide.

## Acknowledgements

I am grateful to the Head of IMMT, Bhubaneswar for allowing me to carry out the summer training in their laboratory. I am thankful to the Head of Department of Ocean Studies and Marine Biology, Pondicherry University to provide me necessary support during the initial sample collection in Andaman Islands.

